# Conserved and lineage-restricted gene regulatory programs modulate developmental cnidocyte specification in *Nematostella vectensis*

**DOI:** 10.1101/2025.05.08.652877

**Authors:** Benjamin Danladi, Layla Al-Shaer, Jamie A. Havrilak, Dylan Faltine-Gonzalez, Kejue Jia, Wyatt Forwood, Timothy Dubuc, Jacob Musser, Michael J Layden

## Abstract

Cnidocytes are a synapomorphy of cnidarians that have evolved a range of morphologies and functions within and across extant species, which makes them an excellent model to investigate how novel cell types emerge and radiate in evolution. One way to gain insight into how cell types evolve is to investigate the gene regulatory networks (GRNs) that pattern them, leading to the identification of shared and divergent programs that likely represent components of the ancestral and evolved phenotypes. Efforts to identify early acting transcription factors in the sea anemone *Nematostella vectensis* revealed that *NvfoxE-like* is a lineage-restricted cnidocyte transcription factor. Expression and functional studies confirmed that *NvfoxE-like* and its targets are found exclusively in cnidocytes. By investigating interactions with regulators of the known cnidogenesis GRN and comparing the targets of known cnidocyte regulators, we identified at least four regulatory programs contributing to the cnidogenic GRN in *Nematostella*. The previously identified cnidocyte regulators *Nvznf845* and *NvpaxA* represent two core pan-cnidarian regulatory programs, while the *NvfoxE-like* dependent program appears to be restricted to *Hexacorallia* or *Actinaria*, where we hypothesize it functions to pattern the unique capsule features present in those lineages. Here, we confirm the hypothesized core conserved transcriptional program, as well as demonstrate how a novel transcription factor was co-opted into cnidogenesis in a subset of cnidarians, thus contributing to the diversification and continued evolution of cnidocytes.

## Introduction

Cnidocytes (stinging cells) are a defining synapomorphy of cnidarians (e.g., anemones, corals, jellyfish, hydra), and they serve critical functions in food acquisition and defense ^1^. There are three main subtypes of cnidocytes: nematocytes (piercing), spirocytes (ensnaring), and ptychocytes (adhesive) ^2^. However, variations in harpoon morphology, venom composition, capsule structure, and function within each of those classes create an incredible diversity of cnidocyte morphologies. There are phylogenetic signatures to some of the variations observed. For example, spirocytes are restricted to the anthozoan lineage, and morphological features related to how the cnidocyte capsule opens to release the harpoon, such as the operculum, apical cap, and apical flaps, are also restricted to specific lineages ^3,4^. Thus, after the emergence of cnidocytes in the stem cnidarian, a radiation occurred, resulting in the variations observed in extant species. The clear novelty of this cell type, coupled with the fact that they radiated across cnidarians, argues that cnidocytes are an excellent model to investigate the origin and evolution of novel cell types in animals. A key question is how the gene regulatory network (GRN) that specifies cnidocytes development first arose and evolved to drive cnidocyte diversification.

Cnidocytes arise from a common pool of progenitor cells that also give rise to neurons and gland cells^5,6^ and express *soxC* and *soxB* family transcription factors ^7^. For example, *sox* genes are expressed in progenitors that give rise to cnidocytes in *Hydra* and jellyfish, and in *Nematostella, NvsoxC and NvsoxB2* are required for cnidocyte development ^7,8^. This suggests that the role of this family of transcription factors is widely conserved across all cnidarians ^9^.

The first indication that progenitor cells become restricted towards the cnidocyte lineage is the expression of the novel transcription factor *Nvznf845,* which is unique to and conserved across the cnidarians ^10^. znf845 and its target paxA are restricted to the cnidocyte lineage in both *Hydra* and *Nematostella* ^10–12^. Further, functional studies in *Nematostella* confirmed that *znf845* is required for cnidocyte development, and that it is downstream of the *sox* genes ^10^ . Together, these data suggest that *znf845* is a pan-cnidarian cnidocyte specification gene.

To date, the only known cnidocyte sub-type regulator is *Nvsox2* ^13^*. Nvsox2* promotes nematocyte development, likely only within the anthozoans, and it is particularly important to suppress the ensnaring spirocyte-like phenotype in small basitrichs^13^. Although it also regulates harpoon length in the large basitrichs ^13^. Thus, there is emerging evidence for a core program that is shared across cnidarians that consists of a *sox → znf845 → paxA* cascade, and lineage restricted mechanisms that pattern newly evolved cnidocyte subtypes, such as *sox2* regulating specification between nematocyte and spirocyte subtypes in anthozoans ^13^.

Here, we investigated the developmental specification of cnidocytes in the actiniarian sea anemone *Nematostella vectensis*. We previously identified early acting neurogenic transcription factors ^14^, and several of them have been shown to regulate cnidogenesis. However, one gene *NvfoxE-like* (formally called *NvfoxD3-like*), had not been described as a candidate cnidogenesis gene. A combination of single-cell mRNA sequencing (scSeq), bulk mRNA sequencing (RNAseq), and functional assays confirmed that *NvfoxE-like* promotes cnidocyte development, and that it is not part of the established core-conserved *sox → znf845 → paxA* cnidocyte specification program. Investigation of target genes identified following the knockdown of *Nvznf845, NvpaxA,* and *NvfoxE-like* revealed that there are likely at least four distinct regulatory programs that simultaneously act to specify cnidocyte fates in *Nematostella,* of which two of them likely reflect core conserved programs. Interestingly, the *NvfoxE-like* gene is independent of the *sox-*mediated cnidogenesis program and many of its targets are found only in hexacoral species or are unique to *Nematostella*. Our findings imply that *NvfoxE-like* plays a unique role in hexacorallian or actiniarian cnidogenesis, which we hypothesize is related to cnidocyte capsule morphology. Collectively, these findings further elaborate the regulatory network of cnidocyte specification in *Nematostella* and provide potential avenues to investigate core-conserved and lineage-specific cnidogenic GRNs that will provide insights about the origin and diversification of this novel cell type.

## Materials and Methods

### Animal care/ spawning

All animals were maintained at 16℃ in ⅓ strength artificial seawater with a salinity of 11-13ppt and a pH of 8.1 - 8.2 and given weekly water changes. To induce spawning, animals were transferred to a 25℃ incubator overnight, with a light on from 11pm onward, and moved to room temperature at 9am. Fertilized embryos were de-gelled in 4% w/v cysteine solution^15^. Embryos used for experiments were cultured at 24℃ and fixed at appropriate experimental timepoints in 4% paraformaldehyde following a previously published protocol ^16^.

### shRNA-mediated gene knockdown

RNAi-mediated gene regulation was utilized to knock down genes of interest. Primers for the shRNAs were designed as previously described (Additional file 2) ^17^, and the shRNA transcription reaction was performed using an AmpliScribe T7-Flash Transcription Kit (Biosearch Technologies, Lucigen). We used a combination of two shRNAs to achieve knockdown for *NvfoxE-like*, and a single shRNA for *NvpaxA*, *Nvznf845*, *NvsoxC*, and *NvsoxB2*. Injection mix of 1 µg/µl shRNA was microinjected into embryos immediately after fertilization and before embryos started cleaving. For all gene knockdown experiments, control embryos were injected with equal concentrations of an established control shRNA ^17^.

### Colorimetric in situ hybridization

Probes for *in situ* hybridization were made by cloning a region of the mRNA using gene-specific primers (Additional file 2), and a transcription reaction template was made from the cloned mRNA region. Digoxigenin RNA labelling mix (Roche 11277073910) was used for probe transcription reactions. *In situ* hybridization was performed using a previously published protocol^18^, and experiments were quantified blindly and imaged with a Nikon DS-Ri2 color camera using the Nikon Element software. Images were processed in Adobe Photoshop 2024.

### Transgenesis and transgene analysis

*Nvncol3::mOrange* transgenic line was obtained from Yehu Moran’s lab ^18^, and the *NvfoxE-like*::*egfp* transgenic line was made using mega-nuclease-mediated transgenesis as previously describe ^19^. We designed primers to clone the promoter region 2kb upstream of the *NvfoxE-like* transcription start site (Additional file 2). Z-stack images of F0 double transgenic embryos fixed 72 hours post fertilization (hpf) were taken using a Zeiss 880 laser scanning confocal microscope, and image processing was done using the Imaris 10.2 software.

### Transcriptomic analysis following gene knockdown

Bulk mRNA sequencing was performed on gastrula-stage embryos following RNAi-mediated gene knockdown by injecting into single-cell stage embryos as described above ^17,20^. Each replicate consisted of ∼1000 injected embryos. Five paired experimental and control replicates were sequenced for all knockdown experiments, except *NvfoxE-like* which had four. RNA was stabilized by placing embryos (28hpf at 24°C) from each replicate in 1 ml of TRI Reagent (Sigma-Aldrich) and then stored at −20°C until total RNA was isolated as previously described ^21,22^. RNA concentration and purity were measured with an Agilent 2100 Bioanalyzer (all samples RIN ≥ 8.0). Library preparation, Illumina NovaSeq 6000 mRNA sequencing (paired-end, 150bp reads), and read quality filtering and trimming were performed by Novogene Inc. as previously described ^23^.

Sequencing data can be retrieved from the National Center for Biotechnology Institute’s Sequence Read Archive (NCBI SRA). The Galaxy web platform was used to map reads to the *Nematostella* Nvec200 genome and conduct differential gene expression analysis using DESeq2 as previously described ^23–25^. A false discovery rate of *p* ≤ 0.05 and log2(FC) of ± 0.585 was used to determine differential gene expression for the *NvfoxE-like* and *NvpaxA* RNAseq, and an adjusted *p* ≤ 0.05 and log2(FC) of ± 0.585 was used for the *Nvznf845* RNASeq data analysis. Overlap between the targets identified in each RNAseq dataset were visualized using the ChiPlot and Biovenn ^26^.

### Single-cell RNA sequencing data analysis

To obtain single-cell libraries, the Chromium 10X genomics protocol was used following the manufacturer’s instructions for a targeted recovery of 10,000 cells (Chromium Next GEM single cell 3’ Reagents Kit v3.1, Dual Index). An Agilent bioanalyzer was used to assess library quality. The Johns Hopkins Single Cell and Transcriptomics Core performed sequencing and pre-processing of samples. Illumina sequencing was performed, and the raw scSeq reads were processed through default pipelines in Cell Ranger 7.0 with alignment to the *Nematostella vectensis* genome ^25^.

Downstream analyses were performed in R/RStudio (version 2022.12.0+353). Low quality cells were identified and filtered out for quality control as follows: cells expressing < 250 or >10000 genes, or cells containing < 500 or > 50000 unique molecular identifiers, and cells containing > 10% mitochondrial counts ^27^. The DoubletFinder function was used to identify and remove doublets ^28^. Further processing and analysis of filtered data was performed using standard Surat pipelines ^29^. runUMAP (Uniform Manifold Approximation and Projection) and DimPlot functions were used to determine and visualize clusters, and the FeaturePlot and DotPlot functions were used to analyze gene expressions.

### Phylogenetic analysis of Fox, Znf845, and PaxA

We identified protein homologs of *N. vectensis* Fox, Znf845, and PaxA using a combination of BLASTp (version 2.15.0) ^30^ and PROST^31^, a protein language model-based approach for remote homology detection. We performed searches against six cnidarian species (*Acropora digitifera*, *Acropora_tenuis*, *Hydra_vulgaris*, *Nematostella vectensis*, *Stylophora pistillata*, and *Xenia sp*) and four model organisms (*Homo sapiens*, *Mus musculus*, *Drosophila melanogaster*, *Caenorhabditis elegans*). BLASTp searches were performed using BLOSUM62^32^ amino acid substitution matrix, with default gap penalty parameters (11 for gap opening and 1 for gap extension). Both BLAST and PROST results were filtered using an e-value cutoff of 1e-6. Multiple sequence alignments were generated using MAFFT (version v7.471) with a maximum iteration setting of 1000 to refine the alignment and using a local pairwise focus to emphasize the conserved regions of the sequences. Phylogenetic trees were then generated with IQ-TREE (version 2.3.6) with default parameters and 1000 bootstrap replicates.

#### Identification of Fox target orthologs expression

We identified orthologs of target genes for the *foxE-like* transcription factor across cnidarians with OrthoFinder (version 3.0.1b1) using default parameters ^33^. Expression patterns were assessed in previously published datasets for the anthozoans *Nematostella vectensis*^34^ , *Stylophora pistillata*^34^ , and *Xenia sp*.^34^ , and the medusozoan *Hydra vulgaris* ^12^. For *Stylophora*, *Xenia*, and *Hydra* we used dataset versions curated by Levy et al., which generated metacell and cell type family assignments, which were used for plotting. For each target gene ortholog, we generated bar plots and violin plots of raw counts in Seurat ^29^ (version 5.1.0) to illustrate the cell type and family-specific expression profiles.

#### Phylogenetic analysis of cnidarian fox genes

Forkhead (Fox) gene sequences were identified from publicly available genomic and transcriptomic resources. Cnidarian gene models were obtained from either the NCBI Genomes resource ^35^ or Ensembl ^36^. Our dataset included 15 cnidarian species spanning anthozoans, scyphozoans, cubozoans, hydrozoans, and staurozoans. To root the phylogeny and provide broader metazoan context, we included outgroup species from sponges, ctenophores, placozoans, flatworms, nematodes, arthropods, chordates, and one mesozoan species. Outgroup taxa included *Amphimedon queenslandica*, *Spongilla lacustris* (Porifera), *Mnemiopsis leidyi* (Ctenophora), *Trichoplax* sp. H1 (Placozoa), *Schmidtea mediterranea* (Platyhelminthes), *Caenorhabditis elegans* (Nematoda), *Drosophila melanogaster* (Arthropoda), *Mus musculus* and *Homo sapiens* (Chordata), and *Rhopalura muelleri* (Mesozoa: Orthonectida).

Candidate Fox genes were identified using BLASTp and tBLASTn searches against genome and transcriptome assemblies using known forkhead protein sequences as queries. For species lacking well-annotated gene models, putative gene regions were predicted using AUGUSTUS ^37^, and manually curated when necessary. The presence of forkhead domains was confirmed using the SMART protein domain prediction tool ^38^.

Full-length protein sequences were aligned using MUSCLE ^39^ within MEGA11 ^40^, using default parameters. The alignment was visually inspected and manually curated to correct potential misalignments, particularly in highly variable or gapped regions. No trimming or domain-specific filtering was applied; the entire protein sequences were retained for phylogenetic analysis.

Phylogenetic reconstruction was conducted using MrBayes v3.2.7 ^41^, on the CIPRES Science Gateway ^42^. Bayesian inference was performed using the WAG amino acid substitution model with a gamma-distributed rate variation across sites. Two independent runs, each with four Markov chains, were executed for 1,000,000 generations, with sampling every 100 generations. The first 25% of samples were discarded as burn-in. Convergence was assessed by confirming that the average standard deviation of split frequencies was below 0.01.

The resulting consensus tree was visualized using FigTree v1.4.4 (http://tree.bio.ed.ac.uk/software/figtree/). Posterior probability values were annotated on internal nodes to indicate clade support. The final tree was used to assess orthologous and paralogous relationships of Fox genes across cnidarian lineages and other metazoan taxa.

### Reciprocal best hit search

We used the NCBI BLASTp tool to identify reciprocal best hits between *Nematostella vectensis* and the hydrozoan *Hydra vulgaris*, several hexacoral species, and a couple of octocoral species. *Nematostella* protein sequences were accessed from the genome assembly hosted by the Stowers Institute of Medical Research (https://genomes.stowers.org/starletseaanemone) ^25^, and sequences for *H. vulgaris* strain AEP were accessed from the genome assembly hosted by the National Human Genome Research Institute (https://research.nhgri.nih.gov/HydraAEP/) ^43^. For comparisons to *hexacorallia*, we used gene models from the marine genomics data repository (reefgenomics.org) ^44^ for the following species: *Acropora digitifera*^45^, *Acropora hemprichii*^46^, *Acropora loripes*^47^, *Acropora tenuis*^48^, *Actinia equina*^49^, *Aiptasia sp.*^50^, *Aiptasia* CC7 (Nagarajan, unpublished), *Porites lutea*^48^. For comparisons to *octocorallia*, we used gene models hosted on NCBI for *Xenia sp.* (RefSeq assembly: GCF_021976095.1)^51^ and *Dendronephthya gigantea* (RefSeq assembly: GCF_004324835.1)^52^. We used the protein sequence for the top BLASTp hit identified for all species surveyed to perform a BLASTp comparison against *Nematostella vectensis*. We considered a gene to be a putative homolog if it returned the same *Nematostella vectensis* gene ID as our gene of interest.

## Results

### *NvfoxE-like* is a novel cnidocyte-specific gene

To identify cnidogenesis regulators, we screened our pool of previously identified early acting neurogenic transcription factors^14^ for expression in cnidocytes by mapping them to a *Nematostella* gastrula-stage scSeq atlas. Five genes (*NvpaxA, Nvsox2, Nvcoup-like1* (aka *NR12* ^10^*), Nvdkk3-like3, and NvfoxE-like (aka NvfoxD3-like)*) were highly expressed in cnidocytes (Figure 1A; 1B). Four of these genes (*NvpaxA, Nvsox2, Nvcoup-like1,* and *Nvdkk3-like3*) are previously identified cnidogenic genes (Figure 1A - B, Figure S2A - F) ^8,10,13,53^. However, *NvfoxE-like (aka foxD3-like)* was highly enriched in cnidocytes and their progenitors (Figure 1B, large asterisk), and had not been previously described as a cnidocyte regulator. Since *NvfoxE-like (aka foxD3-like)* had not been investigated phylogenetically, we first sought to validate its relationship to known *fox* genes. *NvfoxE-like* was formally named *NvfoxD3-like,* but we find that it strongly groups with the foxE gene family rather than the foxD family, which appears to have undergone several gene duplications in early cnidarian evolution (Figure S1, Additional file 1), hence, we refer to this gene as only *NvfoxE-like* onwards.

**Figure 1.**
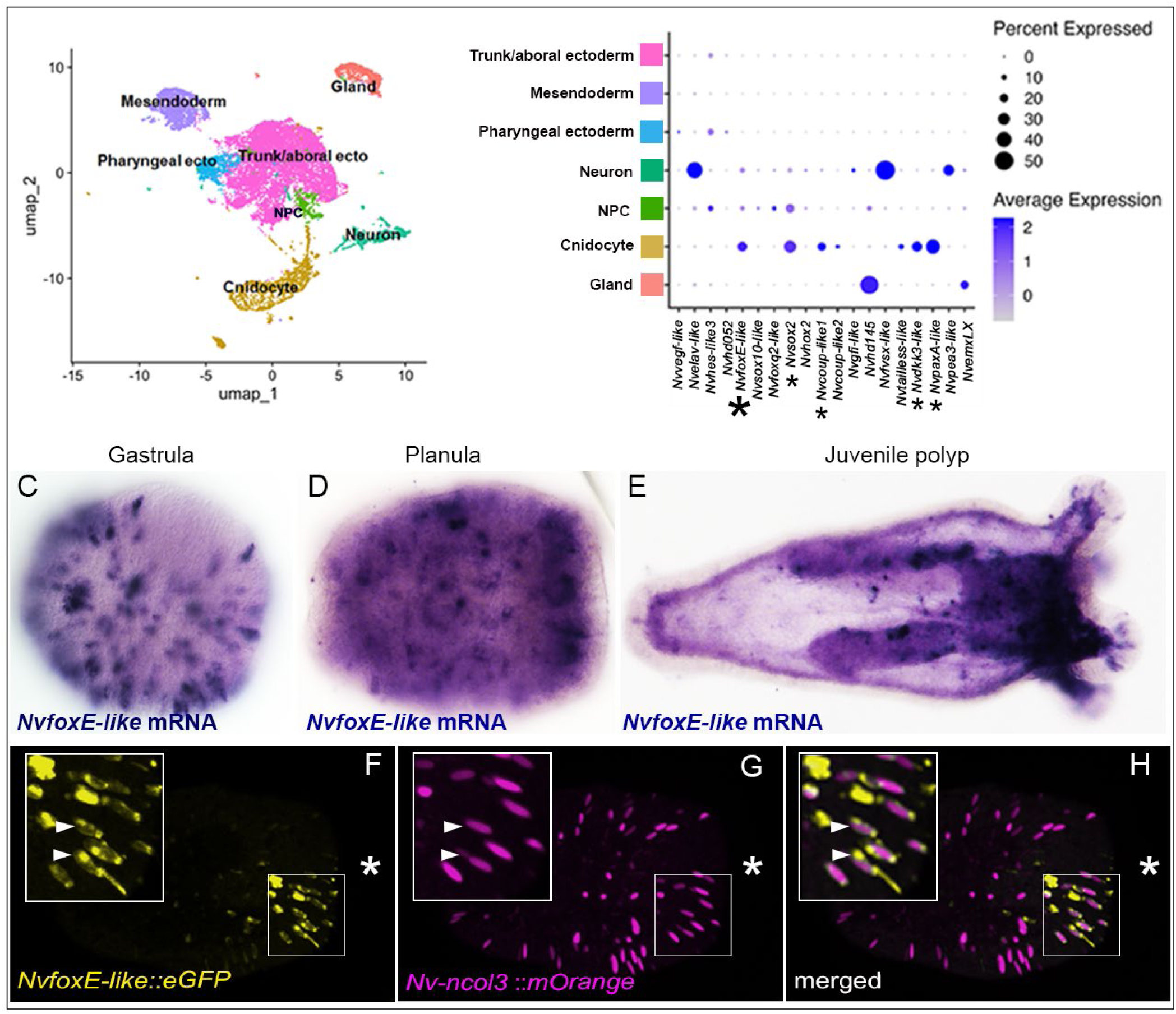
*NvfoxE-like* transcription factors have cnidocyte-enriched expression. (A) UMAP showing annotation of cell clusters from a gastrula stage scSeq atlas. (B) Cellular expression of transcription factors downstream of the MEK signaling pathway. Small asterisks denote transcription factors with known cnidocyte expression and the large asterisk labels *NvfoxE-like*.). (C) *NvfoxE-like* mRNA expression at gastrula), (D) planula, and (E) tentacle-bud stages. (F-H) A double transgenic line of *NvfoxE-like*::eGFP and *Nvncol3*::mOrange showing colocalization (white triangles) in a planula stage embryo. The white asterisk in panels F-G marks the oral end of the embryo

Next, we assessed the expression of *NvfoxE-like*. Using scSeq atlases generated from gastrula through late planula developmental stages, we found that the expression of *NvfoxE-like* was restricted to cnidocytes and their progenitors (Figure 1C, Figure S2 G - I). Additionally, mRNA *in situ* hybridization (ISH) confirmed that *NvfoxE-like* is expressed in a salt and pepper pattern throughout development, as would be expected for a cnidocyte regulator. Its expression is reduced in the body column by late-planula stages when cnidocyte specification is waning but remains strong in tentacles which are constantly undergoing cnidogenesis (Figure 1D - F)^53^. Lastly, mosaic expression of the F0 reporter *NvfoxE-like::egfp* transgene injected into *Nvncol3::morange* cnidocyte reporter lines showed significant overlap between *NvfoxE-like* and *Nvncol3* expressing cells (Figure 1G – I). We conclude that *NvfoxE-like* is expressed in cnidocytes and that it is a putative regulator of cnidocytes in *Nematostella*.

### *NvfoxE-like* functions independent of the conserved *Sox→znf845→paxA* program

Because the progenitor genes *NvsoxC* and *NvsoxB(2)* are known to be the most upstream cnidogenic genes ^8^, their requirement for *NvfoxE-like* expression was tested. Knockdown of *NvsoxC* and *NvsoxB(2)* together or individually results in a loss of cnidocytes and a reduction in expression of *Nvznf845* and *NvpaxA* ^8^. However, loss of either *NvsoxC* or *NvsoxB(2)* had no impact on *NvfoxE-like* expression, and double knockdown of both genes only mildly reduced *NvfoxE-like* expression (figure 2 A - G, figure S3). This is distinctly different from the strong downregulation of *paxA, znf845,* and other cnidocyte genes following the knockdown of either *NvsoxC* or *NvsoxB(2)* (figure S3) ^5^. The lack of an impact on *NvfoxE-like* expression following the loss of the *sox* genes suggests that *NvfoxE-like* is not a major component of the *sox→znf845→paxA* cnidogenic gene regulatory program.

**Figure 2:**
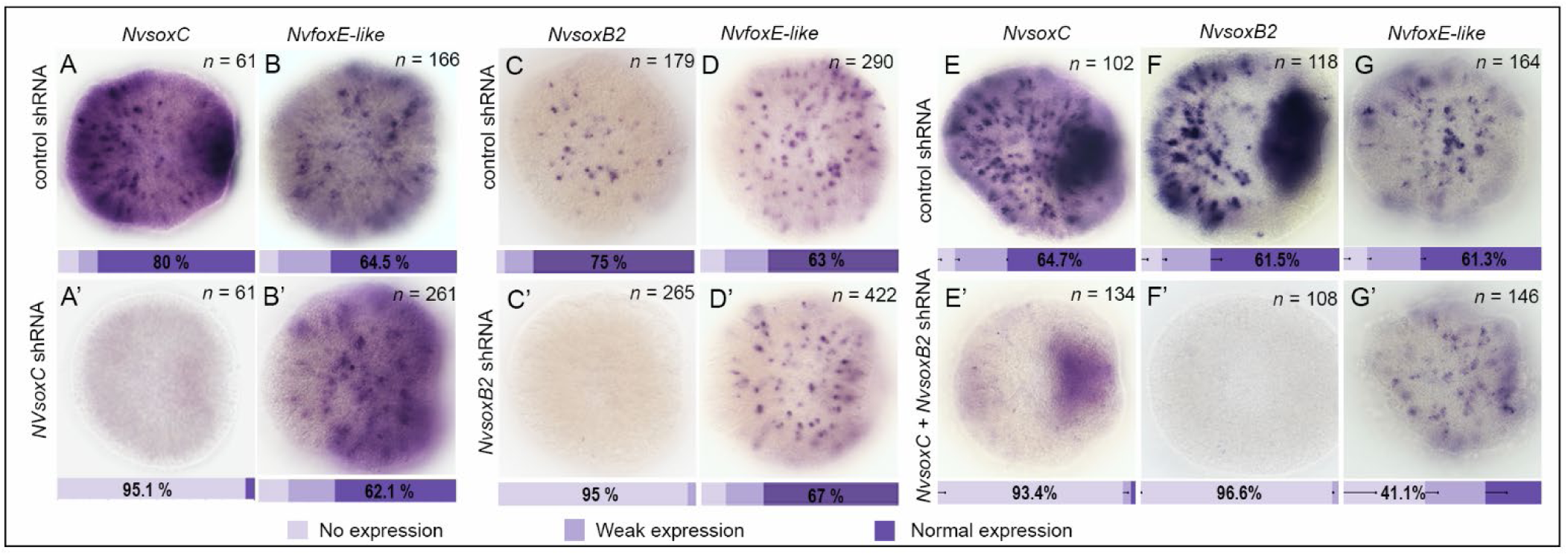
*NvfoxE-like* is not regulated by known upstream cnidogenesis regulators *NvsoxC* or *NvsoxB2*. (A-B’) *NvfoxE-like* expression is not affected by *NvsoxC* knockdown at 30hpf compared to embryos injected with a scramble control shRNA. (C-D’) Similarly, *NvfoxE-like* expression is not affected by *NvsoxB2* knockdown compared to embryos injected with control shRNA.,(E-G’) *NvfoxE- like* expression is only slightly affected by *NvsoxC* + *NvsoxB2* knockdown compared to embryos injected with control shRNA. In all pictures the oral end is to the right. Error bars represent the one-sided standard error of the mean percentage distribution of three technical replicates.

However, our findings do not exclude possible negative interactions of *NvfoxE-like* with *Nvznf845* and *NvpaxA* or vice versa. To determine if *NvfoxE-like* interacts with these known cnidogenic transcription factors, the expression of each gene was screened in different knockdown backgrounds (Figure 3). Knockdown of *Nvznf845* had no measurable effect on *NvfoxE-like* expression (figure 3A - B, Additional file 5), and *Nvznf845* expression was not disrupted following *NvfoxE-like* knockdown (figure 3C - D), which suggests no interaction between the two transcription factors. Similarly, the loss of *NvpaxA* did not significantly change the expression of *NvfoxE-like* and *vice versa* (figure 4E - H). These findings were consistent with RNAseq results, indicating no interaction between *NvfoxE-like* and *NvpaxA* (Additional files 2-3). Collectively, these data argue that *NvfoxE-like* functions in parallel to or redundantly with the known *sox→znf845→paxA* cnidogenic program.

**Figure 3:**
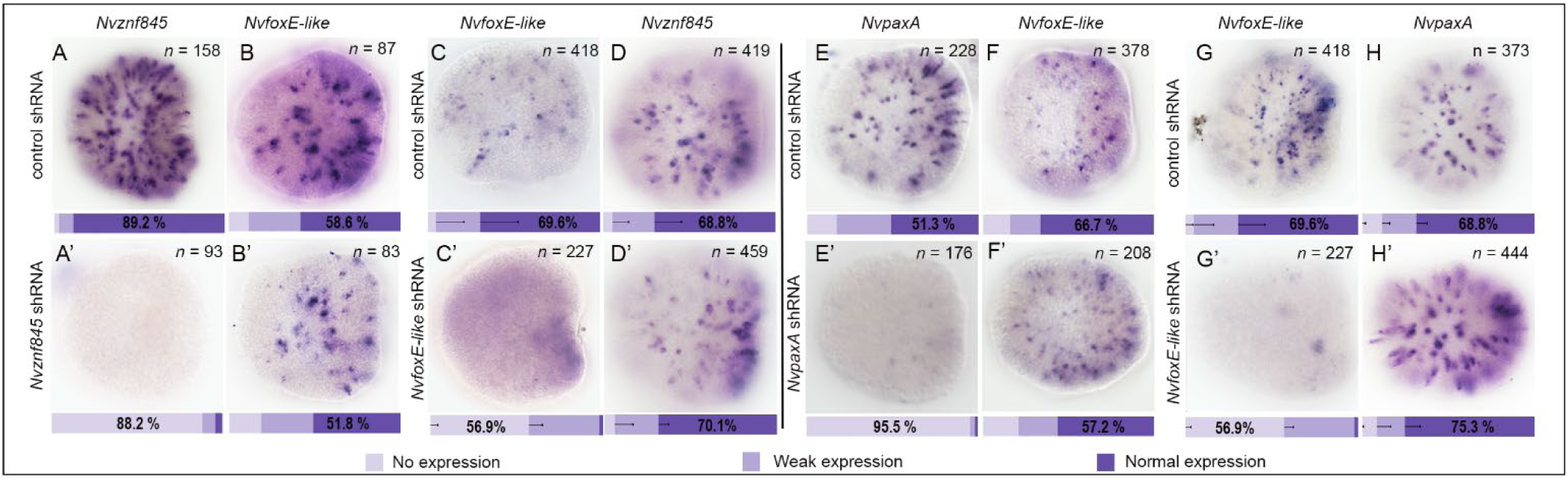
*NvfoxE-like* does not interact with known cnidogenesis genes. (A-B’)*Nvznf845* expression is lost in embryos injected with the znf845 shRNA, but *NvfoxE-like* expression is not affected. (C-D’) *NvfoxE-like* expression is lost in embryos injected with *NvfoxE-like* shRNA, but *Nvznf845* expression is unaffected under the same condition. (E-F’) *NvpaxA* expression is lost in *NvpaxA* shRNA injected embryos, but *NvfoxE-like* expression is unaffected by *NvpaxA* knockdown. (G-H’) *NvfoxE-like* expression is lost in *NvfoxE-like* shRNA injected embryos, but *NvpaxA* expression is unaffected. All embryos are positioned with their oral ends to the right. Error bars represent the one-sided standard error of the mean percentage distribution of two technical replicates.

**Figure 4:**
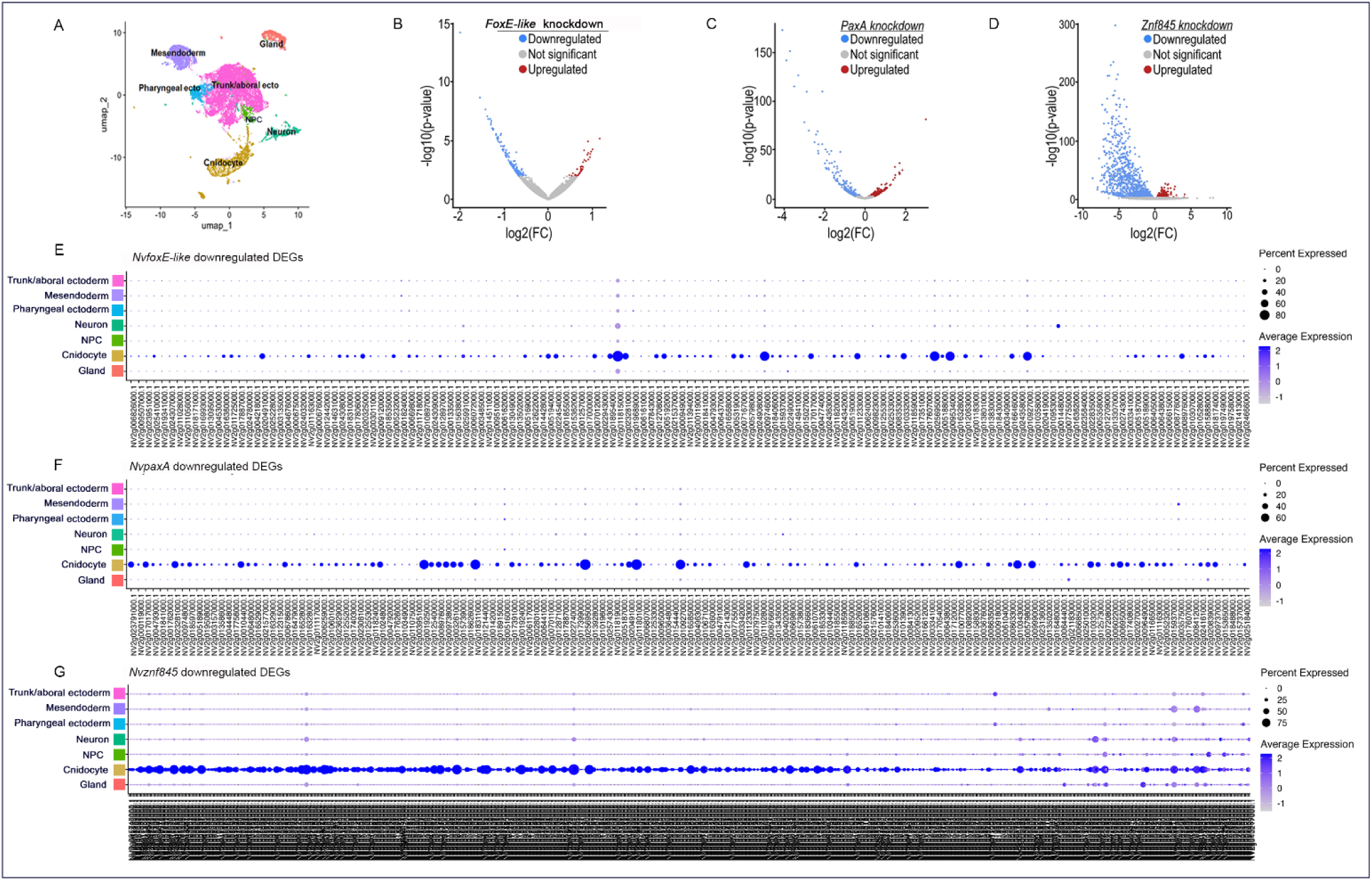
*NvfoxE-like*, *NvpaxA*, and *Nvznf845* downregulated genes are restricted to the cnidocyte cell lineage. (A) A UMAP showing annotated cell clusters in a gastrula-stage scSeq dataset. Volcano plots showing the distribution of genes differentially expressed following shRNA knockdown of (B) *NvfoxE-like*, (C) *NvpaxA*, (D) and *Nvznf845* embryos compared to control embryos. Dot-plots showing the distribution of differentially expressed genes across cell clusters for (E) *NvfoxE-like*, (F) *NvpaxA*, (G) and *Nvznf845 shRNA* injected embryos compared to control shRNA injected embryos.

### Multiple gene regulatory programs contribute to cnidogenesis in *Nematostella vectensis*

To better understand the roles of each gene regulatory program and gain insights about the potential additional programs that promote cnidocyte fate, we performed knockdown of *Nvznf845, NvpaxA,* and *NvfoxE-like* during gastrula stages followed by RNAseq. Targets of each transcription factor were determined by identifying differentially expressed genes (DEGs) observed following shRNA knockdown of each gene relative to stage-matched embryos injected with a scramble shRNA control (Figure 4 B -D). We focus on downregulated DEGs for further analysis to focus on genes that contribute positively to cnidogenesis. RNA-seq identified 958 downregulated DEGs in *Nvznf845* knockdown animals, 154 in *NvpaxA*, and 148 in *NvfoxE-like* (Figure 4B-D, Additional files 2-4). To confirm that these genes likely represent cnidocyte-specific targets of each transcription factor, each gene was mapped to our gastrula stage scSeq atlas. Dot plots revealed strong enrichment in cnidocytes for the downregulated targets of each transcription factor (Figure 4E-G), which is consistent with the known function of *Nvznf845* and *NvpaxA* and further supports our conclusion that *NvfoxE-like* is also a cnidogenic transcription factor.

Because functional studies of *NvfoxE-like* have not been reported, we sought to validate its targets. The list of targets identified by RNAseq was split into terciles and ranked by decreasing adjusted p-value significance. A subset of the downregulated DEGs was selected from each tercile across the whole list of DEGs. We effectively cloned templates for ISH mRNA probes or designed qPCR primers for 13 genes (seven genes, two genes, and four genes for the first, second, and third tercile respectively). The six genes validated by ISH all showed salt and pepper expression in control animals that were decreased following *NvfoxE-like* knockdown (Figure S4). Similarly, the remaining seven genes assessed by qPCR following *NvfoxE-like* knockdown all showed a decrease in expression relative to control. These independent observations of decreased expression in 13 genes following *NvfoxE-like* knockdown using ISH or qPCR recapitulates our RNAseq results and validates our list of *NvfoxE-like* target genes. It should also be noted that the RNAseq results revealed that six out of ten genes previously identified to be highly enriched cnidocyte genes are also targets of *NvfoxE-like* (Figure S5)^18^. In contrast, only one of them is a target of *NvpaxA* (Figure 4) ^18^. Collectively, these data argue that *NvfoxE-like* functions within the cnidocyte GRN in *Nematostella*.

To further characterize how *Nvznf845, NvpaxA,* and *NvfoxE-like* targets fit into the cnidogenic regulatory network, we compared the overlap between the targets of the three transcription factors. In agreement with previously published data, we found that *Nvznf845* regulates *NvpaxA* (Figure 3, Additional Document 2)^10^, supporting the role of *NvpaxA* in cnidogenesis downstream of *Nvznf845.* One hundred and eighteen (118, (96%)) of the genes downregulated by *NvpaxA* knockdown are also downregulated by *Nvznf845* knockdown (Figure 5), but *NvpaxA* has far fewer targets than *Nvznf845,* indicating that *Nvznf845* also promotes some aspects of cnidogenesis independently of *NvpaxA* (Figure 5; Figure 7). One hundred and fifteen (115, (78%)) of the *NvfoxE-like* targets are shared with the *Nvznf845/NvpaxA* program, and 22% (31 genes) are exclusive *NvfoxE-like* targets. These observations imply there are at least four unique programs that promote cnidocyte development (Figure 6) (a *znf845-dependent* program, a *znf845 → paxA* program, a *znf845 | foxE-like* program, and a *foxE-like* program). Next, genes unique to each program were mapped to the *Nematostella* scSeq cnidocyte subset containing data from developmental timepoints and adult samples^54^. Interestingly, the *NvfoxE-like* targets are restricted to maturing and matured cnidocytes and do not show any specific subtype pattern (Figure 7B). This suggests that the *NvfoxE-like* dependent program (Figure 6 green arrows) plays a role in cnidocyte maturation. Its unique targets, *soxA* and *cnido-fos* are known markers of mature cnidocytes ^8^. Further support for the role of *NvfoxE-like* in cnidocyte maturation is that shared *znf845* and *foxE-like* targets also show a strong signature in maturing cnidocytes, but *znf845* and *znf845/paxA* targets are not found in mature or maturing cnidocytes (Figure 7). As expected for a broad and early upstream cnidocyte fate specification gene, unique targets of *Nvznf845* are expressed in all cnidocyte cell populations. Still, surprisingly, not many targets were found in matured cnidocytes (Figure 7C). This distribution of targets appears similar in the program receiving inputs from shared *NvfoxE-like* and *Nvznf845* targets (Figure 7D). This may indicate that both programs (Figure 6 black and purple arrows) broadly act along all stages of cnidocyte differentiation and across all cnidocyte subtypes. Finally, the *Nvznf845*-dependent program acting through *NvpaxA* (znf845 → PaxA, Figure 6, red arrows) appears to promote nematocyte rather than other cnidocyte subtypes (Figure 7E). However, at the gastrula stage, spirocytes are largely absent ^8^, hence, we cannot definitively claim that the paxA regulatory program promotes nematocytes only, it could just be a poor representation of spirocyte-specific genes in our gastrula-stage RNAseq data. There was no clear evidence that either the *NvpaxA* or *NvfoxE-like* pathways regulate cnidocyte subtype patterning. The hypothetical roles for these programs will require further testing, but our analysis of newly discovered and existing cnidocyte genes strongly hints that the regulatory networks driving cnidogenesis branch downstream of *Nvznf845* and potentially include another parallel program that receives input from *NvfoxE-like,* independent of *Nvznf845*.

**Figure 5:**
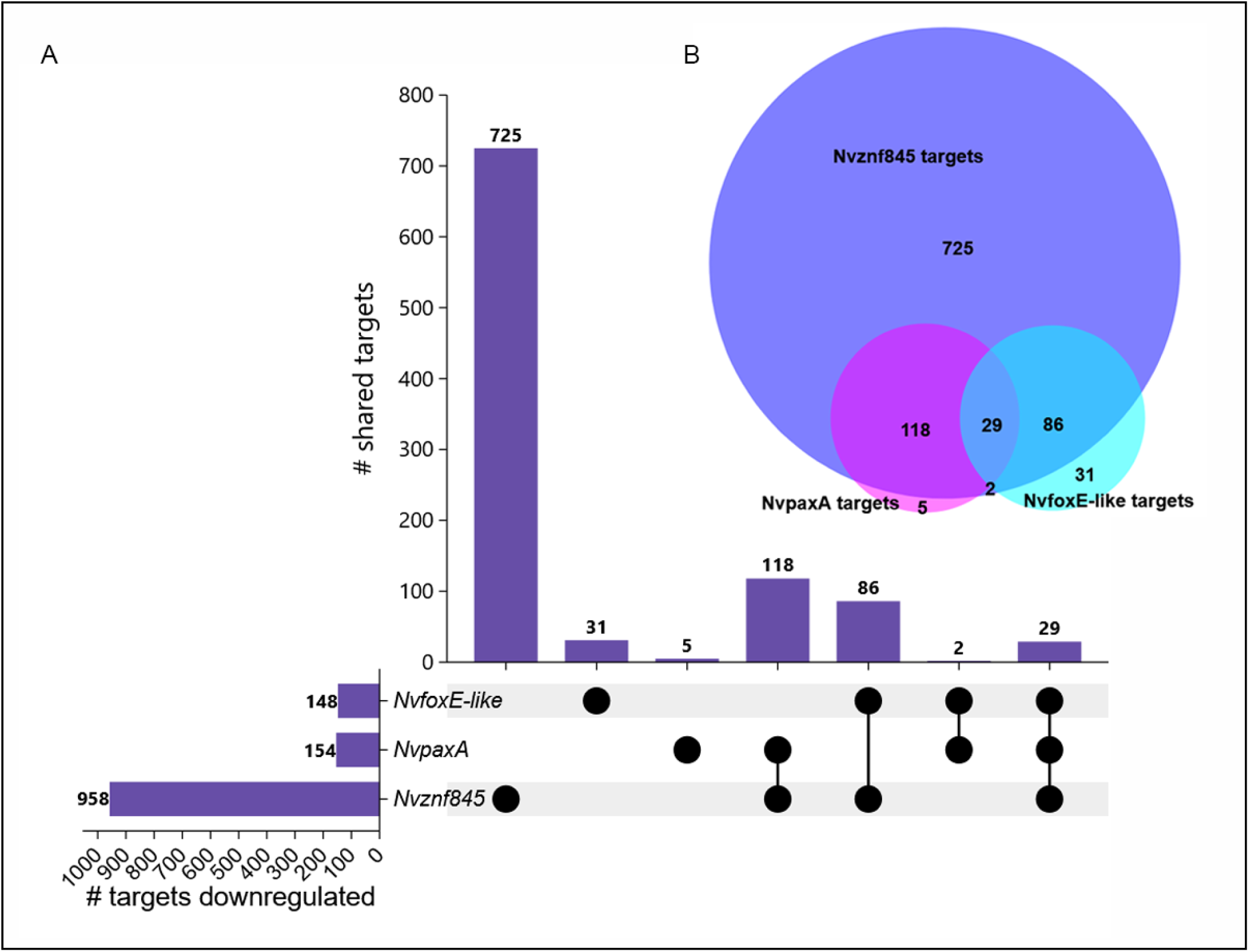
*NvfoxE-like* and *Nvznf845* regulate overlapping and unique targets. (A) Upset plot describing the overlap between Nvznf845, *NvpaxA*, and *NvfoxE-like* targets. (B) Venn diagram of overlap between *Nvznf845, NvpaxA*, and *NvfoxE-like* targets.

**Figure 6:**
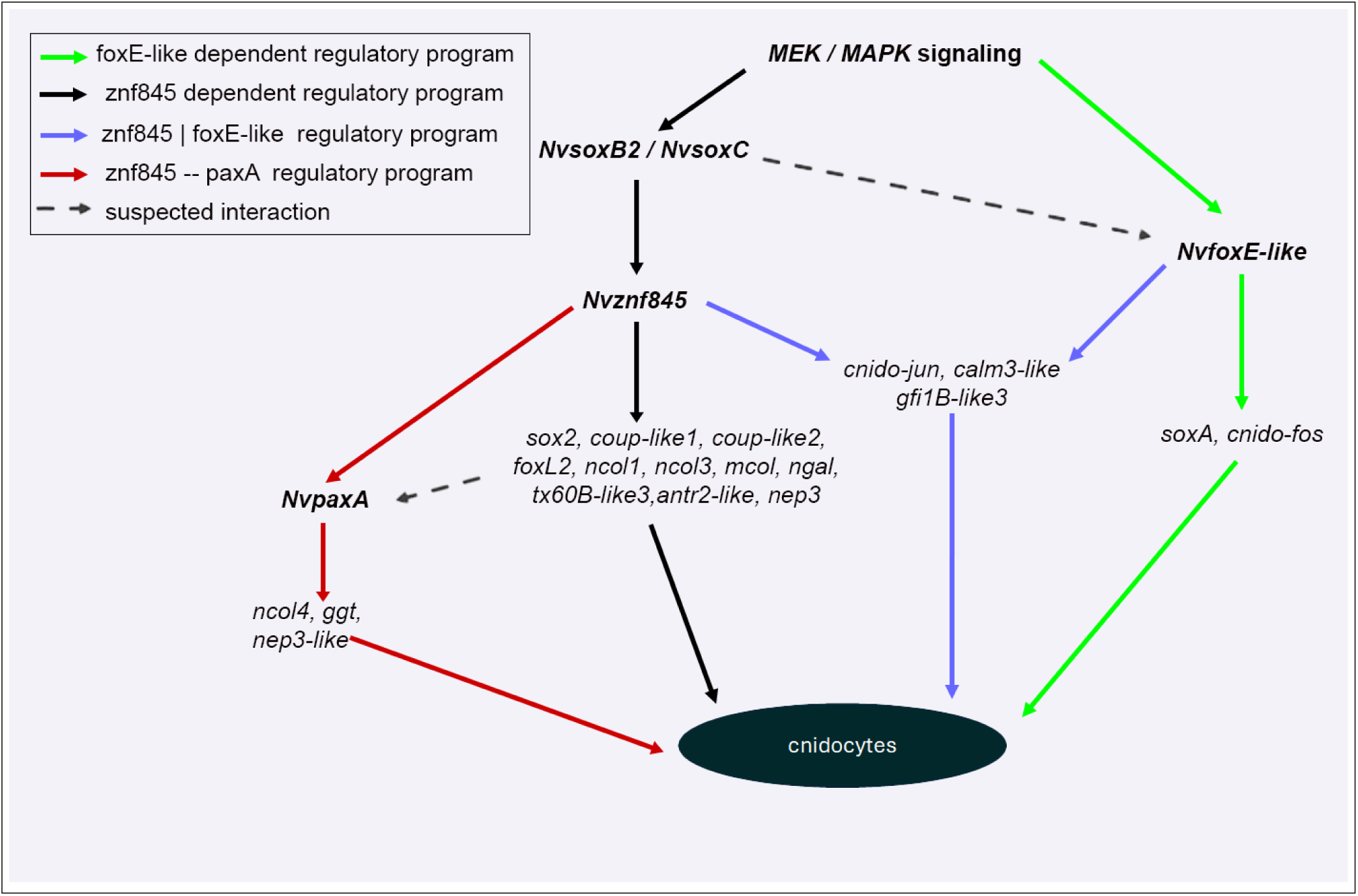
Proposed GRN for cnidocyte specification during development. At least four programs contribute to cnidogenesis. A znf845 independent program (black arrows), a znf845 → PaxA program (red arrows), a znf845 | foxE-like program (purple arrows), and a. foxE-like independent program (green arrows). Arrows with dashed lines indicate suspected interactions

**Figure 7:**
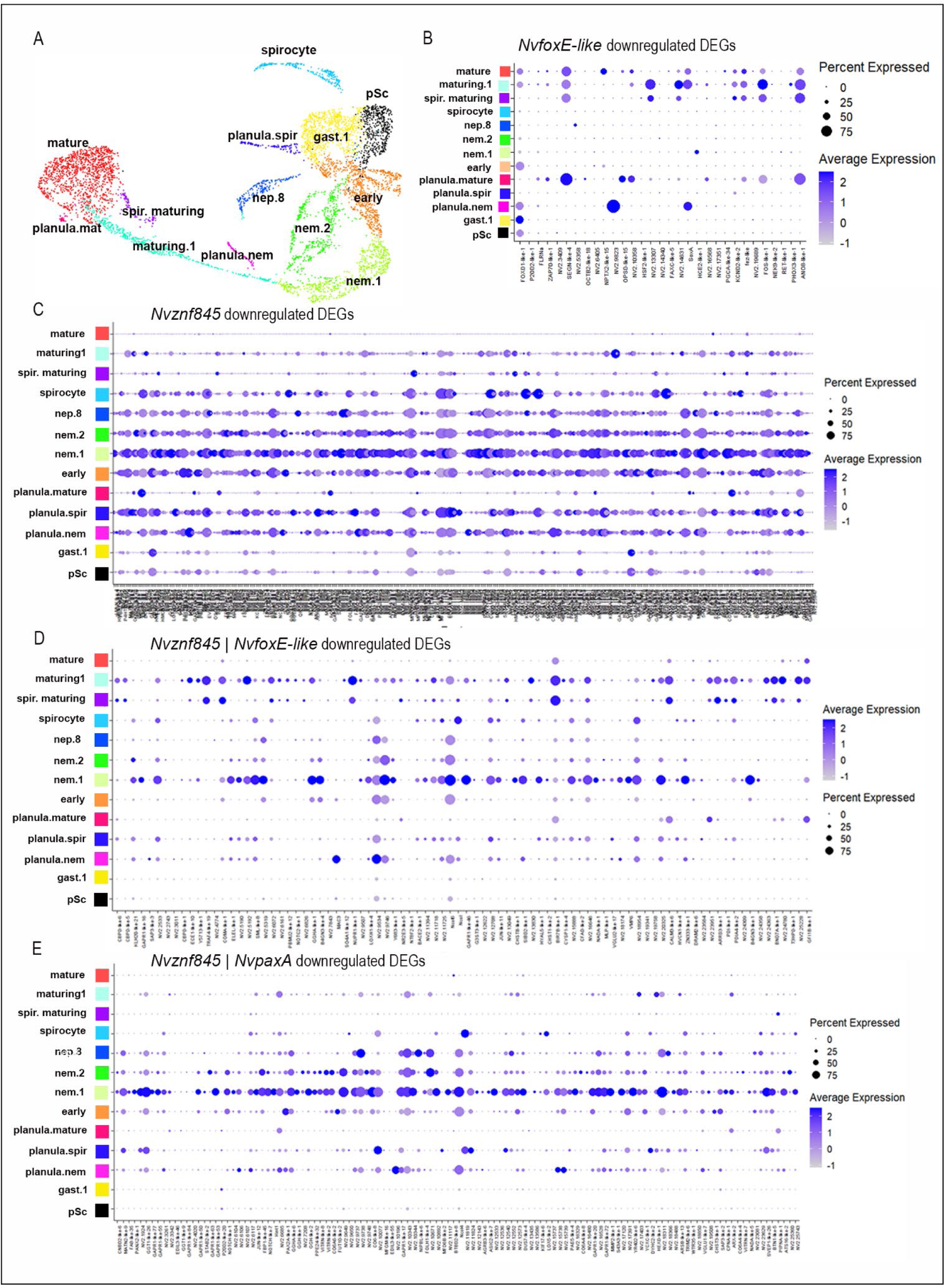
*NvfoxE-like* expression is not restricted to a cnidocyte subtype. (A) UMAP of a *Nematostella* cnidocyte cell subset showing annotated cell states adapted from^54^. (B) Distribution of *NvfoxE-like* unique targets across the cnidocyte cell states (C) Distribution of *Nvznf845* targets across cnidocyte cell states (D) Distribution of *NvfoxE-like* and *Nvznf845* shared targets across cnidocyte cell states (E) Distribution of *NvpaxA* and *Nvznf845* shared targets across cnidocyte cell states.

It is worth noting that mapping previously identified cnidocyte genes to our hypothetical programs provided further support for the predicted programs (Table S1). For example, *Nvznf845* regulates most of the previously identified genes, including capsule structural proteins (*ncol1, ncol3, mcol, ncol4, ngal*)^54^, venom proteins (Tx60B-like3, ANTR2-like, NEP3)^8^, and key regulatory transcription factors (*sox2, coup-like1, coup-like2, foxL2*) (Figure 5) ^10,13^ . *NvpaxA* did not regulate many of the previously described cnidocyte genes, with just one minicollagen gene (*ncol4*) identified as a *NvpaxA* target (Table S1, Figure 6). The fact that many known cnidocyte genes are downstream of *Nvznf845,* but not *NvpaxA,* further supports a role for *znf845* that is independent of the *paxA* program (Figure 6, black arrows). *cnido-jun, gfi1b-like,* and *calm3-like* are targets of both *NvfoxE-like* and *Nvznf845*, indicating that the list of 86 overlapping *NvfoxE-like* and *Nvznf845* targets (Figure 5) likely represents the presence of a program that requires inputs from both transcription factors (Figure 6, purple arrows).

*NvfoxE-like* regulates two transcription factors (*soxA* and *cnido-fos*) independent of either *NvpaxA* or *Nvznf845*, supporting the idea that the 31 unique *NvfoxE-like* targets represent the presence of at least one *znf845* independent program in *Nematostella* (Figure 5, green arrows). Interestingly, the observation that 10/42 of the previously published cnidocyte genes we collated (Table S1) are not regulated by either *Nvznf845*, *NvpaxA*, or *NvfoxE-like*, implies that there may be an unknown cnidogenic program that functions independently of *znf845* in *Nematostella*.

In summary, our data suggest that at least four programs contribute to the cnidogenesis GRN in *Nematostella*. First, a program where *Nvznf845* acts independently of other known regulators of cnidogenesis (Figure 6, znf845 program - black arrows). Second is a program where *Nvznf845* acts upstream of *NvpaxA* (Figure 6, Znf845 → PaxA program, red arrows), and third is a program where *Nvznf845* acts in synergy with *NvfoxE-like* (Figure 6, Znf845 | foxE-like program-purple arrows). Finally, a program where *NvfoxE-like* acts independent of other known regulators of cnidogenesis (Figure 6, foxE-like program - green arrows).

### *NvfoxE-like* dependent cnidogenic program is likely restricted to the *Hexacorallia* or the *Actinaria*

Phylogenetic analyses were performed to determine if the foxE-like dependent program is likely broad across cnidaria. Analysis of proteins containing the DNA-binding domain of the forkhead box across cnidarian clades revealed that *NvfoxE-like* is found only in hexacorals (Figure 8A, Additional file 8), suggesting that it may be a *Hexacorallia*-unique innovation. Next, the *NvfoxE-like* targets were analyzed. Reciprocal blast of *NvfoxE-like* down-regulated DEGs against the *Hydra vulgaris* (Medusozoan) genome^19^ yielded only 31 (21%) putative homologs (Figure 8B, Additional file 6). This data hints that the *NvfoxE-like* program may not be present in the medusozoan lineage. Next, we assessed two classes within the anthozoan lineage; 80% of the top 20 *NvfoxE-like* DEGs had a putative homolog in hexacorals, but only 30% had a putative homolog in the octocorals assessed (Figure 8C, Additional file 7). Similarly, 15 of the 31 unique *NvfoxE-like* targets (48.4%) had putative homologs in hexacorallian genome databases, and only 4 (12.9%) had putative homologs in octocorallian genome databases on NCBI (Figure 8D, Additional file 7). Notably, we did not find reciprocal blast hits for the known cnidocyte genes *soxA* and *cnido-fos* in the octocoral *Xenia sp.*^51^, and *Dendronephthya gigantea* ^52^ genomes (Table S1, Additional file 7). It should be noted that of the genes assessed, ∼30% of the *NvfoxE-like* targets are unique to *Nematostella* (Additional file 7, Additional file 9), not all hexacoralian species assessed had a clear *NvfoxE* homolog, and the *NvfoxE-like* homolog in *Stylophora* was not detected in the single cell atlas. Thus, it is not clear if *NvfoxE-like* homolog expression in cnidocyte lineages is widespread across the hexacorals. We cannot rule out that *NvfoxE-like* functions broadly in the hexacorals, but we currently favor the hypothesis that *NvfoxE-like’s* role in cnidogenesis is restricted to the *Actinaria*.

**Figure 8:**
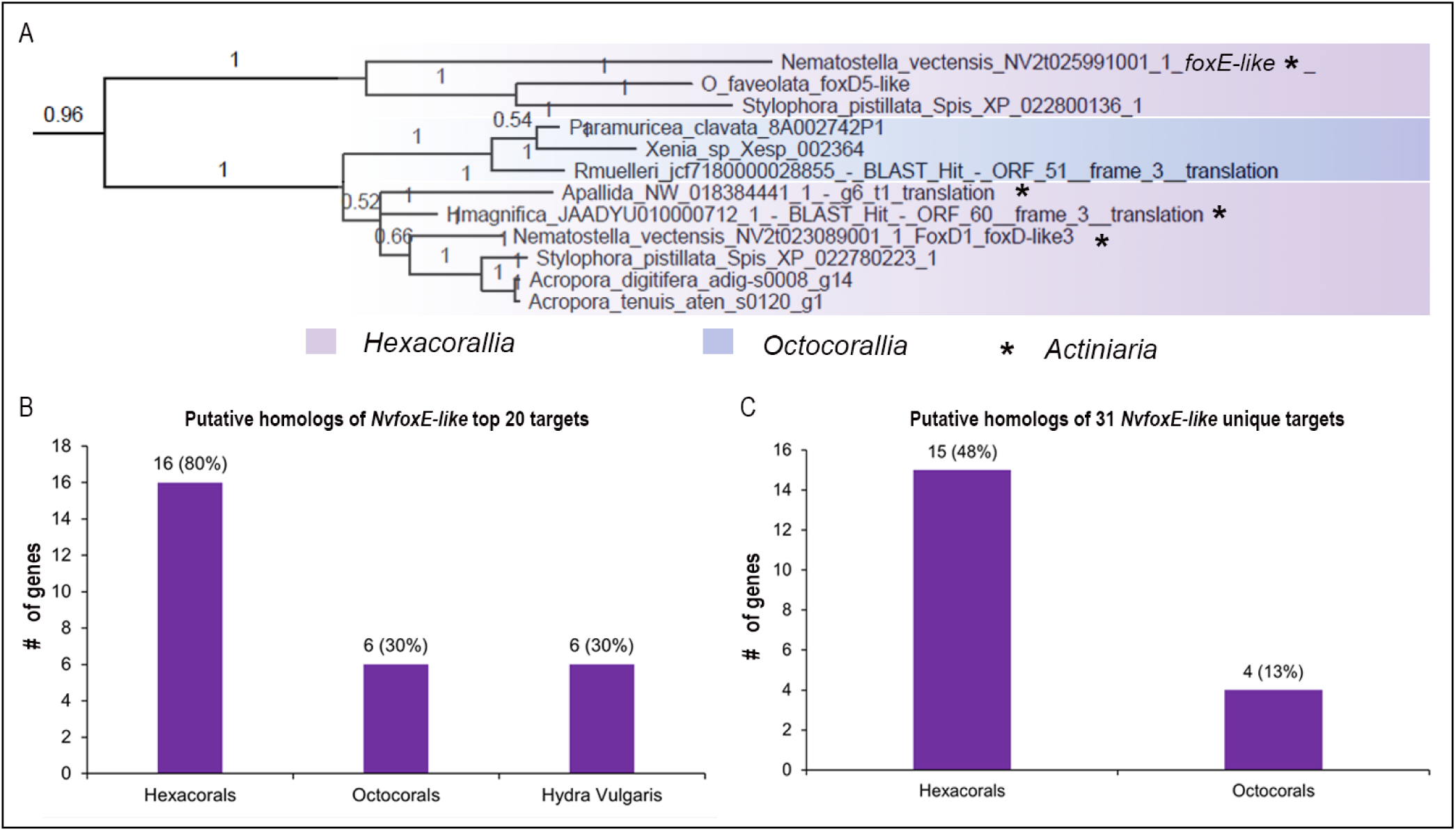
*NvfoxE-like* is restricted to the Hexacorallia lineage. (A) Phylogenetic position of *NvfoxE-like* with other proteins containing the forkhead DNA binding domain across cnidaria. (B) Putative homologs of *NvfoxE-like* targets in the medusozoan *Hydra vulgaris and* select hexacoral and octocoral genome databases. (C) Putative homologs of the 31 *NvfoxE-like* unique targets in select hexacoral and octocoral genome databases.

## Discussion

### Confirmation of proposed core programs, and identification of lineage restricted regulatory program

Our findings provide insights into the GRN regulating cnidocyte development in *Nematostella* and provide the initial framework to elaborate the regulatory networks that define the shared and lineage specific gene regulatory programs that led to the emergence and diversification of the novel cnidocyte cell type.

Previous studies argued that *znf845* and *paxA* homologs broadly regulate cnidocyte fates across all cnidarians ^10–12^. The broad distribution of *znf845* and *paxA* across all cnidarians, and the robust expression of genes in the *Nvznf845* and the *NvpaxA* program across all cnidocyte cell states and developmental time-points strongly suggest that this regulatory program may represent the core pan cnidarian program that led to emergence of cnidocyte in the stem cnidarian. In *Nematostella,* the *Nvznf845* program (Figure 6, black arrows) split into at least three distinct programs. In addition to the known *NvpaxA* dependent pathway, there is clear evidence that *Nvznf845* has a program that is independent of *NvpaxA,* because most of its targets are not shared with *NvpaxA* (Figure 5B, 7C). A third program functions in parallel with or in coordination with *NvfoxE-like* to regulate a common set of target genes. *Nvsox2* regulates small basitrich cnidocyte fate, and harpoon morphology in large basitrichs, and it has been shown that *NvpaxA* acts downstream of *Nvsox2* ^13^, this may represent another paxA dependent program where *Nvznf845* regulates *NvpaxA* via *Nvsox2* (Figure 9). RNAseq analysis of *Nvsox2* target will provide better resolution of this pathway.

**Figure 9:**
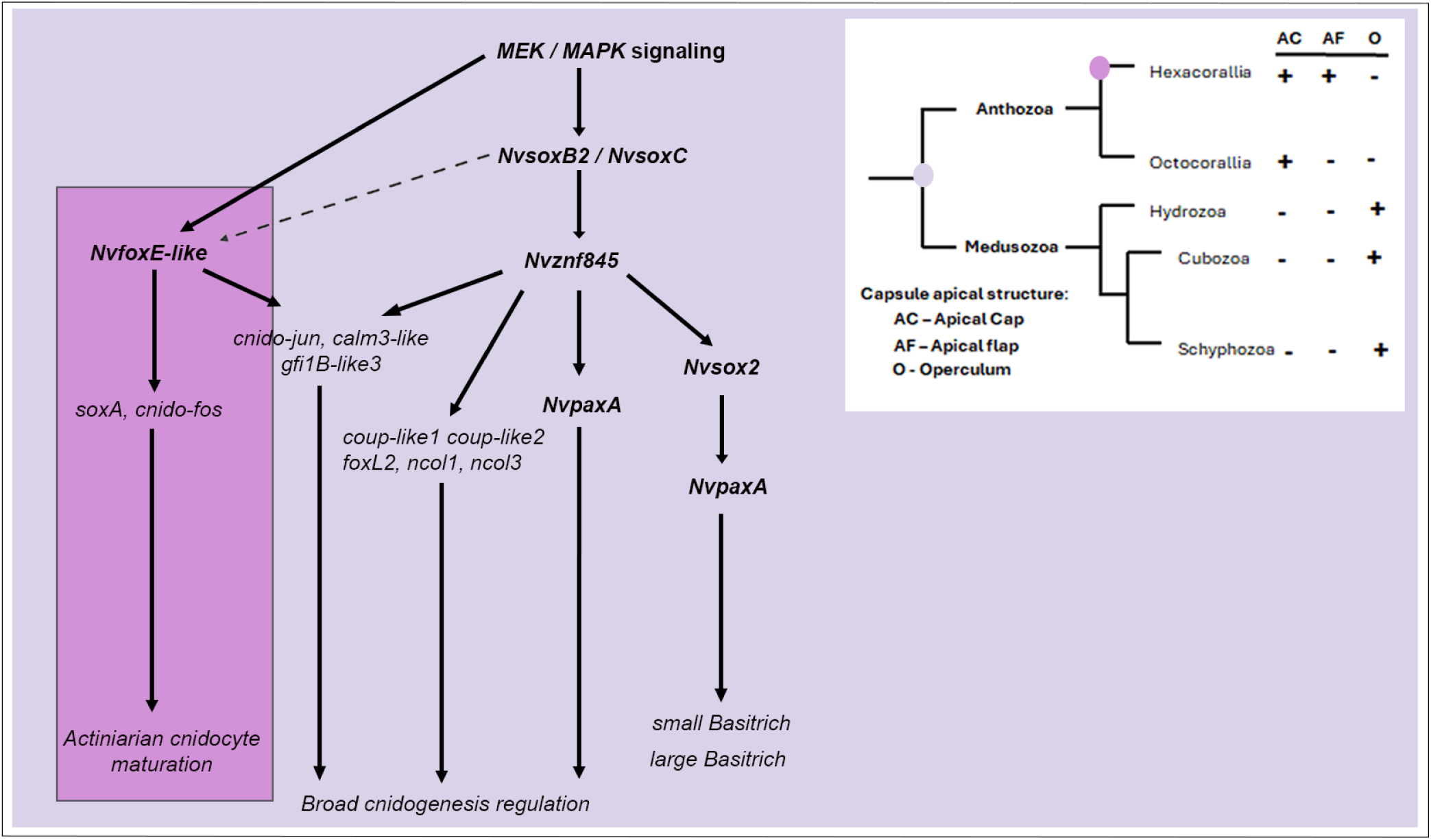
Hypothesized cnidogenesis GRN in *Nematostella vectensis*. Hypothetical roles of program contributing to cnidogenesis GRN and the inset is a phylogeny showing distribution of the apical structure of the cnidocyte capsule, showing apical flap is restricted to hexacorals (actiniarians)^3^

To get better resolution of all these programs, detailed analysis of cnidogenesis at all stages of cnidocyte development including adults stage samples will help resolve the role of each of the regulatory programs in cnidocyte specification.

The *foxE-like* program (Figure 6, green arrows) appears to be restricted to the hexacorallian or actiniarian lineage, *NvfoxE-like* and most of its top 20 downregulated DEGs and unique targets are restricted to *Hexacorallia* or *Nematostella* (an actiniarian) (Figure 8). The *NvfoxE-like* targets are enriched in the matured, and maturing cnidocytes (Figure 7B), and include *Cnido-fos* and *soxA* previously shown to mark matured cnidocytes ^8^. These genes do not have a clear homolog in octocoral or other cnidarians. This further supports our hypothesis that the foxE-like program is restricted to hexacorallians or actiniarians and suggests that this program functions to promote features of the matured hexacoral or actiniarians cnidocyte. To date the most significant difference in mature cnidocytes of the hexacoral or actiniarians lineages is the presence of the apical flaps in *Actiniaria*. Medusozoans possess an operculum, whereas anthozoans have an apical cap and within the anthozoans the *Actiniaria* (which includes *Nematostella*) have some cnidocytes that possess apical flaps (Figure 9)^3^ . Previous studies in *Nematostella* found examples of both the apical cap and apical flap depending on cnidocyte subtype^13^. Given the distribution of *foxE-like* and its targets outside of the *Actinaria*, our best hypothesis is that *NvfoxE-like* functions to regulate the apical morphology of actiniarian cnidocytes. Attempts to recover a mutant for these *NvfoxE-like* by our lab were unsuccessful, making it difficult for us to test this hypothesis. However, to date, it is the only distinguishing feature we are aware of that is broadly distributed across hexacorallian cnidocyte subtypes, that is not found in other lineages. Regardless, it is clear that we identified several genes that are known to be expressed in *Nematostella* cnidocytes that are uniquely downstream of *NvfoxE-like*.

Taken together, our data provide evidence for multiple cnidogenic programs that function both broadly and in lineage restricted contexts to regulate cnidocyte specification. While the core gene regulatory program drove the emergence of cnidocyte in the stem cnidarian, emergence of regulators such as *NvfoxE-like* drove the radiation of cnidocyte diversity across cnidarian clade. Further studies into the detailed regulatory networks and their impacts on cnidocyte fate will shed light on how this unique cell type emerged and diversified during animal evolution.

## Acknowledgements

We would like to acknowledge Leslie Babonis for the feedback that led to conducting the RNAseq on *Nvznf845* and *NvpaxA* and Yehu Moran for sharing the *Nvncol3::mOrange* line with us. We also acknowledge Whitney B Leach and Marc Meynadier for their feedback and help with troubleshooting experiments. Lastly, we thank Fabian Rentzsch, Simon Sprecher, and members of their labs for feedback on the project at various stages of our investigation. This work was funded by NSF CAREER 1942777 and NIH R01 GM127615.

## Author’s contributions

BD: designed and performed experiments, analyzed data, wrote the first draft of the manuscript. LA: performed bulk RNA sequencing experiments and analysis, blindly quantified *in situ* hybridization experiments, injected embryos for knockdown experiments, and performed the reciprocal blast search. JAH: performed and analyzed scSeq experiments, injected embryos for knockdown experiments, and performed ISH. DFG: Performed *Nv*s*oxB2* knockdown qPCR experiment. WF: performed reciprocal blast search. TD: performed the Fox genes phylogenetic analysis. KL and JM: performed phylogenetic and gene homology analysis. MJL: conceived the study, designed the experiments, and wrote the manuscript. All authors read and approved of the final manuscript.

## Materials and correspondence

Michael J Layden

## Supplementary figures

**Figure S1:**
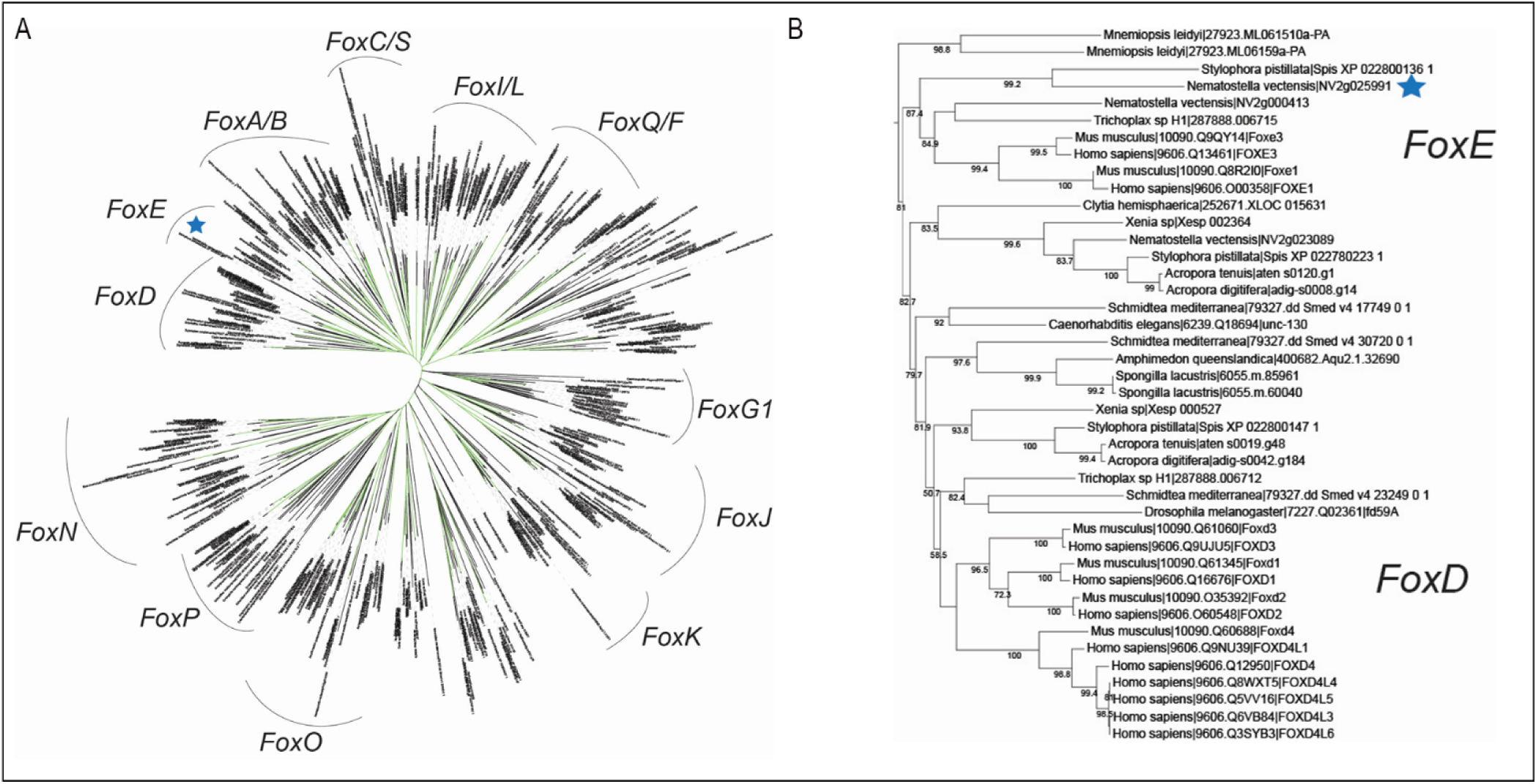
*NvfoxD3-like* is closely related to the foxE family. (A) Gene tree from phylogenetic analysis of forkhead gene, blue star indicates the location of our gene of interest. (B) zoomed-in gene tree, shows that our gene of interest is closely related to the foxE family not the foxD, hence we rename the gene *NvfoxE-like*

**Figure S2:**
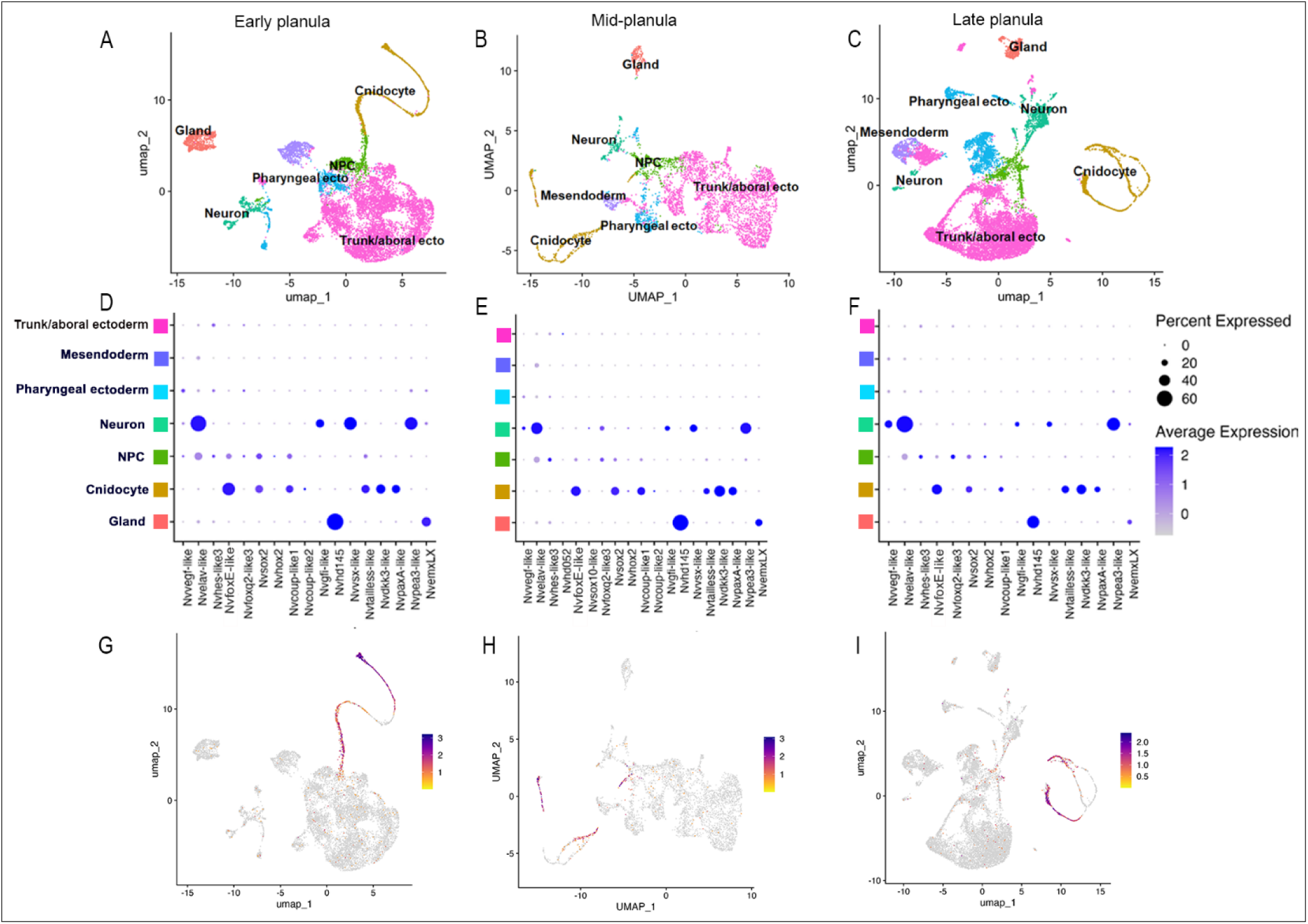
*NvfoxE-like* is expressed in cnidocyte lineages throughout development. Umap showing annotation of different cell types from ScRNA-seq data at early planula (A), mid-planula (B), and late-planula (C) stages. Dot-plot showing MEK signaling downstream transcription factors expressed in the cnidocyte cell population at early planula (D), mid-planula (E), and late-planula (F) developmental stages. Umap showing cell populations expressing *NvfoxE-like* at early planula (G), mid-planula (H), and late-planula (I) developmental stages.

**Figure S3:**
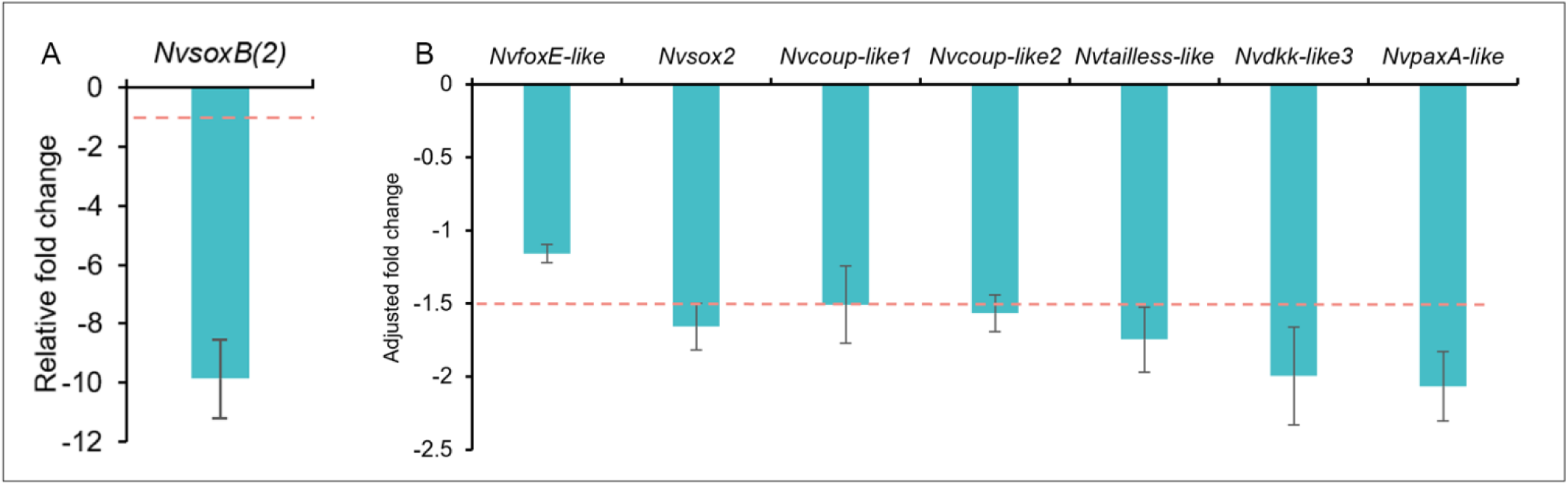
*NvfoxE-like* is not regulated by the upstream cnidogenesis regulator NvsoxB(2). (A, B) qPCR analysis of *NxsoxB(2*) knockdown in gastrula stage embryos. (A) *NvsoxB(2)* expression is significantly reduced in *NvsoxB(2)* shRNA-injected animals, demonstrating an effective knockdown phenotype. (B) Analysis of the expression of downstream MEK signaling factors by qPCR shows that *NvfoxE-like* expression is not affected by the knockdown of *NvsoxB(2)*.

**Figure S4:**
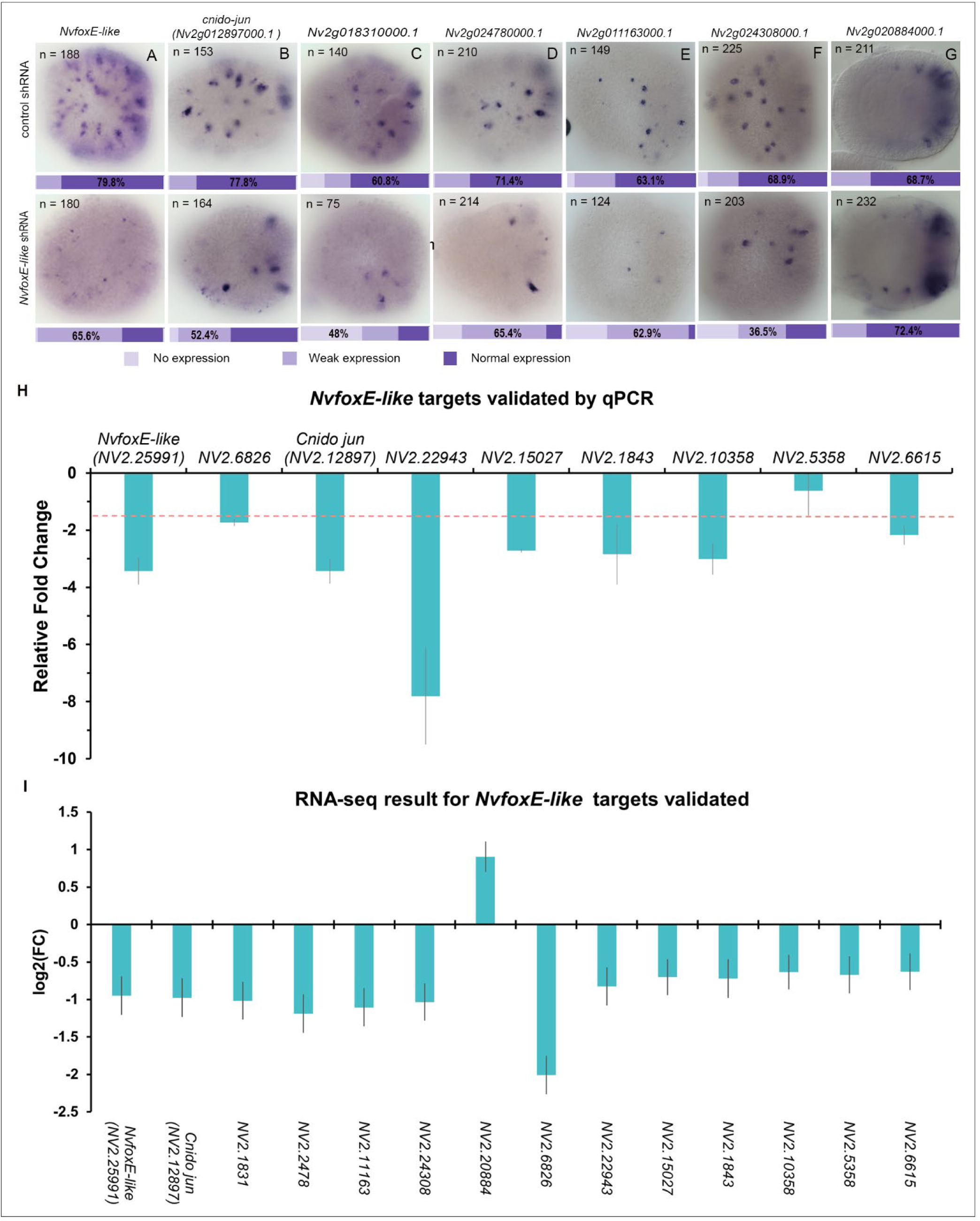
ISH and qPCR validate *NvfoxE-like* RNA-seq data. (A - G) ISH of genes identified by bulk RNA-seq to be differentially expressed by *NvfoxE-like* knockdown recapitulates RNA-seq result, qPCR result of genes identified by bulk RNA-seq to be differentially expressed by *NvfoxE-like* knockdown recapitulates RNA-seq result (H), RNA-seq result of targets validated by ISH and qPCR(I). qPCR experiment was done in triplicate, representing different biological samples

**Figure S5:**
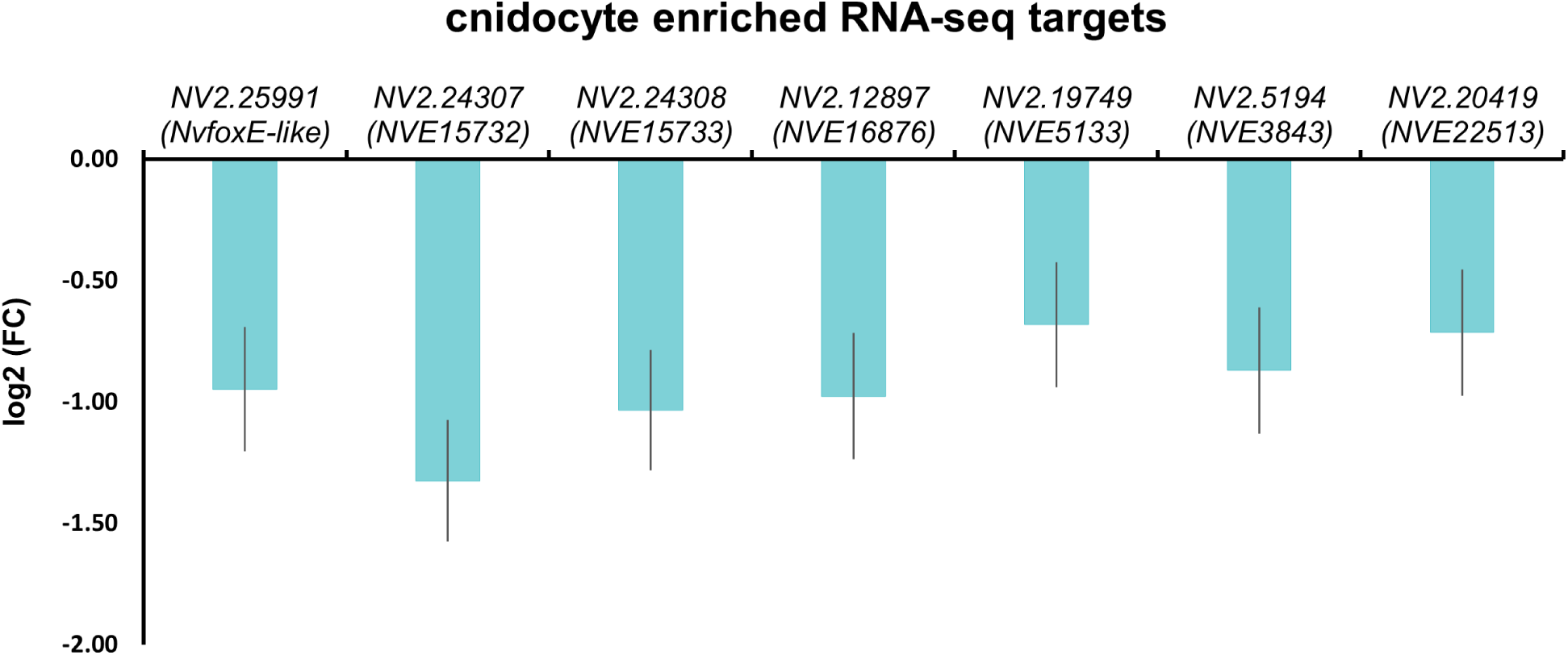
*NvfoxE-like* regulates known cnidocyte genes. RNA-seq data showing *NvfoxE-like* regulates targets previously identified to enriched in cnidocytes ^11^.

**Table S1:** Distribution of published cnidocyte genes regulated by *NvfoxE-like*, *NvpaxA*, and N*vznf845*. “+” indicates the corresponding transcription factor regulates the cnidocyte gene.

| Gene name | NV2 ID | NVE ID | *znf845* | *foxE-like* | *paxA* |
| --- | --- | --- | --- | --- | --- |
| *znf845* | NV2.17877 | NVE16303 |  |  |  |
| *sox2* | NV2.18482 | NVE2102 | + |  |  |
| *ncol1* | NV2.12621 | NVE4226 | + |  |  |
| *ncol3* | NV2.10686 | NVE9976 | + |  |  |
| *mcol1* | NV2.12623 | NVE4225 | + |  |  |
| *Ngal* | NV2.5200 | NVE3845 | + |  |  |
| *coup-like1* | NV2.23247 | NVE21135 | + |  |  |
| *coup-like2* | NV2.1477 | NVE17305 | + |  |  |
| *dkk3-like3* | NV2.10134 | NVE13825 | + |  |  |
| *NVE26200* | NV2.23901 | NVE26200 | + |  |  |
| *NVE5730* | NV2.21271 | NVE5729 | + |  |  |
| *NVE13546* | NV2.22446 | NVE13546 | + |  |  |
| *foxL2* | NV2.14459 | NVE1324 | + |  |  |
| *TX60B-like3* | NV2.11971 | NVE21250 | + |  |  |
| *ANTR2-like* | NV2.21578 | NVE16011 | + |  |  |
| *NEP3* | NV2.20490 |  | + |  |  |
| *myc4* |  | NVE20964 | + |  |  |
| *PRDM6-like* |  | NVE25286 | + |  |  |
| *cnido-jun* | NV2.12897 | NVE16876 | + | + |  |
| *NVE15732* | NV2.24307 | NVE15732 | + | + |  |
| *NVE15733* | NV2.24308 | NVE15733 | + | + |  |
| *NVE3843* | NV2.5194 | NVE3843 | + | + |  |
| *CALM3-like* | NV2.20419 | NVE22513 | + | + |  |
| *GFI1B-like3* | NV2.25410 | NVE16640 | + | + |  |
| *Ncol* | NV2.11820 | NVE18955 | + | + |  |
| *foxE-like* | NV2.25991 |  |  |  |  |
| *SoxA* | NV2.15027 | NVE426 |  | + |  |
| *cnido-fos* | NV2.19749 | NVE5133 |  | + |  |
| *ncol4* | NV2.11801 | NVE18974 | + |  | + |
| *ggt* | NV2.1925 | NVE20501 | + |  | + |
| *paxA* | NV2.6786 | NVE25857 | + |  |  |
| *NEP3-like* | NV2.20491 | NVE22462 | + | + | + |
| *NVE17236* | NV2.10392 | NVE17236 |  |  |  |
| *ATF2-like1* | NV2.11500 | NVE20585 |  |  |  |
| *myc5* | NV2.18928 | NVE21040 |  |  |  |
| *Nkx2.2D* | NV2.11134 | NVE10557 |  |  |  |
| *foxA* | NV2.11441 | NVE20630 |  |  |  |
| *Six1-2* | NV2.11090 | NVE9850 |  |  |  |
| *Tx60B-like5* | NV2.12610 | NVE4235 |  |  |  |
| *NEP8* | NV2.19484 | NVE15921 |  |  |  |
| *Max-like* |  | NVE12845 |  |  |  |
| *PRDM13* |  | NVE7535 |  |  |  |

**Additional file 1**: Phylogenetic analysis of foxE-like

**Additional file 2**: List of primers used.

**Additional file 3:** List of DEGs in foxE-like knockdown RNA-seq

**Additional file 4:** List of DEGs in paxA knockdown RNA-seq

**Additional file 5:** List of DEGs in znf845 knockdown RNA-seq

**Additional file 6:** Reciprocal blast hits of foxD-like2 targets in hydra vulgaris genome

**Additional file 7:** Reciprocal blast hits of top 20 foxE-like targets and foxE-like unique targets in the genome of other anthozoans

**Additional file 8:** Phylogenetic tree for genes with the conserved forkhead box (fox) DNA binding domain across cnidarians

